# DNA repair by Rad52 liquid droplets

**DOI:** 10.1101/768119

**Authors:** Roxanne Oshidari, Richard Huang, Maryam Medghalchi, Elizabeth Y.W. Tse, Nasser Ashgriz, Hyun O. Lee, Haley Wyatt, Karim Mekhail

## Abstract

Cellular processes are influenced by liquid phase separation, but its role in DNA repair is unclear. Here, we show that in *Saccharomyces cerevisiae*, Rad52 DNA repair proteins at different DNA damage sites assemble liquid droplets that fuse into a repair centre droplet. This larger droplet concentrates tubulin and projects short aster-like microtubule filaments, which tether the droplet to longer microtubule filaments mediating the mobilization of damaged DNA to the nuclear periphery for repair.

---

Eukaryotic genomes are dynamic structures and are non-randomly arranged within the cell nucleus, which is defined by an envelope perforated with nuclear pore complexes (NPCs)^1^. Genome dynamics allow cells to repair DNA double-strand breaks (DSBs), which are highly toxic DNA lesions that trigger the DNA damage checkpoint^2^. Specifically, the movement of DSBs allows them to escape repair-repressive heterochromatin domains, search for homologous sequences or localize to repair-conducive NPCs^3–15^.

The *de novo* assembly of intranuclear filaments, onto which DSBs are transported by motor proteins, promotes DSB escape from heterochromatin or movement to NPCs^6,16–18^. In *S. cerevisiae* cells with a single DSB, the Kinesin-14 motor proteins Kar3 and Cik1 associate with the break site and are required for its capture by long DNA damage-inducible intranuclear microtubule filaments (DIMs), which emanate from the microtubule-organizing centre (MTOC)^6,16^. The break is then directionally mobilized by Kinesin-14 onto a DIM and moved away from the MTOC to NPCs for repair^16^. Similarly, in cells treated with carcinogens such as methyl methanesulfonate (MMS), damaged DNA, identified by the presence of the Rad52 DNA repair protein, moves along DIMs to NPCs, where the focus later dissolves, marking repair completion^16^. In flies, a similar actin/myosin-based mechanism moves DSBs for repair^8,18^. Importantly, in a given cell, carcinogens can trigger several DSBs that co-localize and create a DNA repair centre, which is enriched in Rad52 in yeast but remains poorly understood across eukaryotes^17,19^. The forces driving DSB clustering, whether such forces crosstalk with nuclear filaments, and how clustering promotes genome stability remain unclear.

Therefore, we used a yeast system for the fluorescence-based visualization of DSB-indicating Rad52, α-tubulin Tub1, and NPC-indicating Nup49 protein^16^ (Supplementary Fig. 1a-b). Cells treated with MMS exhibited Rad52/RPA-positive DSBs (Supplementary Fig. 1c). MMS induced one DIM in cells containing a single large and bright Rad52 focus (Fig. 1a-b). DIMs emanated from the MTOC and efficiently captured the large Rad52 focus, as expected (Fig. 1c)^16^. In contrast, the MTOC of cells containing more than one Rad52 focus tended to exhibit several shorter microtubule filaments (denoted petite DIMs or pti-DIMs) that failed to capture damaged DNA (Fig. 1a-c). Thus, cells with several DSB-indicating Rad52 foci exhibit several pti-DIMs, which, in contrast to the DIM in cells with one large Rad52 focus, fail to capture the Rad52 foci.

**Fig. 1.**
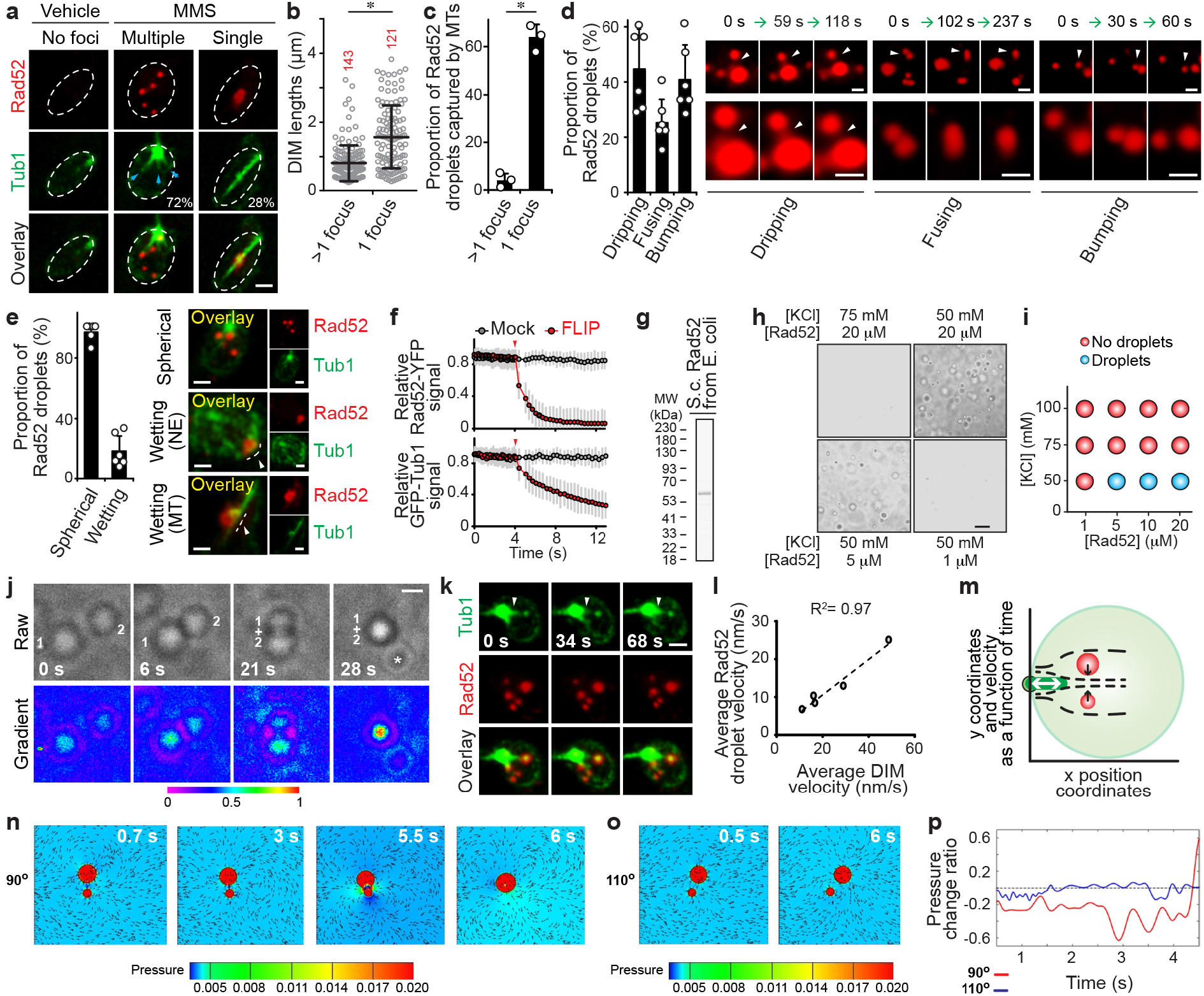
DSB-indicating Rad52 forms liquid droplets whose fusion can be explained by flow-generating short nuclear microtubules. **a-c**, Live-cell microscopy shows that nuclei with >1 Rad52 foci (**a**) exhibit shorter microtubule filaments (**b**) that cannot capture the foci (**c**) (n=3). In (**a**), shown are % of cells with given DIM species and pti-DIM-indicating blue arrowheads. **d-e**, Rad52 foci exhibit liquid droplet behaviour (*n*=6). **f**, Rad52 foci dynamically exchange constituents with the nucleoplasm (Rad52 *n*=30, Tub1 *n*=12). **g-i**, Rad52 forms liquid droplets *in vitro* in a salt/protein concentration-dependent manner. **j**, Rad52 droplets fuse *in vitro*. Asterisk, separate droplet entering the imaging frame. **k-l**, pti-DIM extension/shortening (**k**, white arrowhead) correlates with Rad52 droplet velocity (**k, l**). **m-p**, CFD-based modelling of pti-DIM motion shows that it can generate flow (**m**) that drives Rad52 droplet fusion when pti-DIM velocity is applied at ~90° (**n**) but not at 110° (**o**) relative to the axis connecting the centre of the droplets. Fusion is driven by an inter-droplet decrease in pressure that is shown in gradients (**n-o**) and quantifications (**p**). Scale bars, 1 μm (**a-e, k**), 10 μm (**h**), 5 μm (**j**). Quantifications represent the mean ± s.d.; \**P* < 0.0001 in Mann-Whitney U (**b**) and χ^2^ (**c**) tests.

Strikingly, in cells harbouring one or more Rad52 foci, the foci exhibited liquid-like properties^20–22^. First, in pti-DIM-positive cells, Rad52 foci that were induced using the genotoxic agents MMS or zeocin engaged in dripping, fusion, or bumping encounters with each other (Fig. 1d; Supplementary Fig. 1d; Supplementary Movies 1-3). Second, in DIM-positive cells, the Rad52 focus exhibited wetting behaviour as it flattened against the nuclear envelope or the DIM (Fig. 1e). Third, Rad52 foci were abrogated by the liquid droplet disruptor 1,6-Hexanediol, which did not disrupt the overall nuclear localization of Rad52 (Supplementary Fig. 1e)^2^. Lastly, Rad52 foci quickly lost signal during fluorescence loss in photobleaching (FLIP), confirming that the foci constituents are liquid-liquid phase-separated but undergo exchange with the surrounding nucleoplasm (Fig. 1f; Supplementary Fig. 1f; Supplementary Movie 4). The data indicate that Rad52 foci exhibit liquid-like properties *in vivo*.

*S. cerevisiae* Rad52 purified from *Escherichia coli* phase separated from buffer and formed liquid droplets at low salt concentrations (Fig. 1g-i)^23,24^. These droplets were spherical, often fused with each other, and were disrupted by 1,6-Hexanediol (Fig. 1j; Supplementary Movie 5; Supplementary Fig. 2a). Consistent with its liquid droplet-forming capacity, Rad52 is predicted to exhibit a high level of intrinsic disorder (Supplementary Fig. 2b). In fact, a Rad52 mutant (*Δ307*) lacking a portion of the disordered domain failed to phase separate *in vitro* (Supplementary Fig. 2c-d). Importantly, upon DNA damage induction in cells expressing *Δ307*, the percentage of cells with repair foci decreases from ~55% to ~5%, the remaining foci-positive cells also exhibit a higher number of smaller foci, and the whole cell population is hypersensitive to DNA damage^25^. In addition, in strains expressing *Δ307* but not wild-type Rad52, sensitivity to MMS can be partly rescued following the overexpression of Rad51^25^. This indicates that *Δ307* is defective in focus formation *in vivo* and fails to phase separate *in vitro*, but has the ability to promote Rad51 loading and strand exchange *in vivo*. Furthermore, within our experimental conditions, addition of the heterotrimeric repair factor RPA in complex with ssDNA did not alter Rad52 phase separation (Supplementary Fig. 2e-f)^26^. While these data show that Rad52 has an intrinsic ability to assemble liquid droplets, *in vitro* conditions likely do not fully recapitulate the *in vivo* environment in which Rad52 phase separates.

We then asked whether 1,6-Hexanediol, which represses Rad52 droplets *in vivo* (Supplementary Fig.1e), hyper-induces the DNA damage checkpoint, which is indicated by the phosphorylation of Rad53 (CHK2 tumour suppressor in mammals)^2^. Only in MMS-treated cells, 1,6- Hexanediol hyper-induced Rad53 phosphorylation, and this without altering cellular Rad52 levels (Supplementary Fig. 3a-b). Thus, 1,6-Hexanediol hinders genome stability only upon DNA damage induction by disrupting Rad52 phase separation or potentially other factors in the cell.

*In vivo*, a single engineered DSB induces one Rad52 focus and one long DIM but no pti-DIMs, suggesting their potential role in DSB clustering^16^. Indeed, in MMS-treated cells, pti-DIMs engaged in extension-shortening cycles with velocities correlating with Rad52 droplet velocities (Fig. 1k-l; Supplementary Movies 6-7). Computational fluid dynamics (CFD) simulations can reveal whether and how a velocity-induced flow drives the fusion of two liquid droplets^20^. Indeed, CFD simulations incorporating parameters observed *in vivo* revealed that pti-DIM dynamics can generate flows that lower the pressure between Rad52 droplets, driving their fusion (Fig. 1m-p; Supplementary Fig. 4a-b; Supplementary Movies 8-9). Fusion occurred only at droplet viscosities ≤ 0.005 Pa s and was very efficient when the pti-DIM velocity was applied at a 90° angle to the axis connecting two droplets. Therefore, the pti-DIMs may be creating nucleoplasmic flow dynamics that drive Rad52 droplet fusion and genome stability.

Consistent with this notion, disruption of pti-DIMs upon deletion of the Tub3 α-Tubulin isoform decreased Rad52 droplet velocities, preventing droplet clustering and hyper-inducing the DNA damage checkpoint (Fig. 2a-f; Supplementary Fig. 5a-b). Thus, pti-DIMs promote Rad52 droplet clustering and genome stability.

**Fig. 2.**
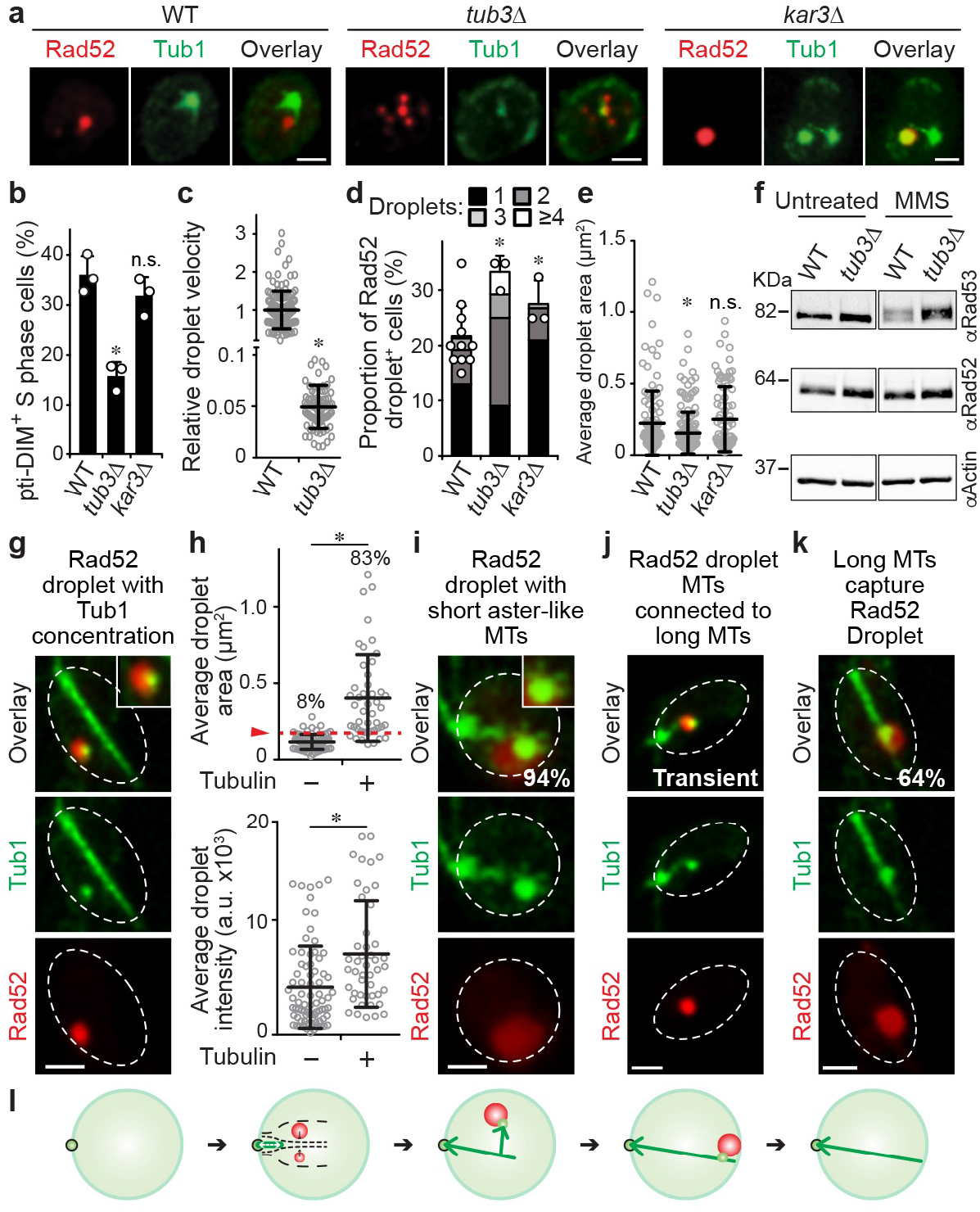
pti-DIMs promote genome-stabilizing clustering of Rad52 droplets and the formation of repair centres that concentrate tubulin and facilitate capture. **a-f**, Live-cell microscopy shows that the pti-DIM-compromising *tub3Δ* (**a-b**) lowers droplet velocity (**c**), increases droplet numbers (**d**), and decreases droplet size (**e**), while western blotting shows Rad53 hyper-phosphorylation (**f**). *kar3Δ* served as control. **g-h**, Live-cell confocal microscopy and quantification showing that Rad52 droplets exceeding 0.2 μm^2^ in size (red line) have a 10-fold higher chance of concentrating tubulin (*n*=6, 168 droplets). **i-k**, Tubulin foci inside Rad52 droplets project aster-DIMs (**i**) that reach the long DIM (**j**) before stable Rad52 droplet capture by the DIM (**k**). Frequency of events is shown in white. MTs, microtubules. **l**, Proposed model. Quantifications represent the mean ± s.d.; *P ≤ 0.0004 in χ^2^ (**b,d**) and Mann-Whitney U (**c,e,h**) tests. Scale bars, 1 μm.

We then asked if the clustering of Rad52-indicated DSBs also promotes their capture by the long DIM, which then guides DSBs to the nuclear periphery. Strikingly, after rounds of Rad52 droplet fusion, when the droplet area reached the threshold size of 0.2 μm^2^, these droplets internally concentrated tubulin into a focus that partially protruded from the droplets (Fig. 2g-h). Subsequently, short aster-like DIMs (aster-DIMs) emerged from the tubulin focus inside the Rad52 repair centre droplet (Fig. 2i). One aster-like DIM became dominant and transiently associated with the long DIM (Fig. 2j). The repair centre droplet then moved along the DIM to the nuclear periphery (Fig. 2k; Supplementary Movie 10), where the Rad52 focus disappeared upon repair completion^16^. Thus, upon the fusion of small Rad52 droplets into a repair centre droplet, the droplet concentrates tubulin and projects aster-like microtubule filaments, which mediate repair centre capture by DIMs for perinuclear targeting and repair.

We found that the cooperation between Rad52 liquid droplets, various nuclear filaments, and nucleoplasmic flow can drive the assembly and function of the DNA repair centre (Fig. 2l; Supplementary Fig. 6). Future work should explore the signals regulating Rad52 droplets. Importantly, a recent study suggested that LLPS of the mammalian DNA repair protein 53BP1 coordinates DNA damage responses at break sites with global p53-dependent gene activation and cell fate decisions^27^. In addition, other proteins have been observed to become liquid-like at sites of DNA damage, suggesting that coordination of the phase separation of several factors at DNA breaks may ensure global genome stability^22,24,28^. We note that another study examining irreparable DSBs did not observe a requirement for Rad52 or filaments in the clustering of such DSBs^29^. Therefore, cooperation between DSB clustering and nuclear filaments in the promotion of DSB mobility likely only occurs in the context of repairable DSB systems. Our results may also point to novel therapeutic approaches, as microtubule-dependent DSB dynamics can drive chromosome translocations and thereby alter the sensitivity of cancer cells to therapy^7^. Overall, we have deciphered repair centre assembly and function, expanded the repertoire of biological liquid droplets, and uncovered hidden dimensions of genome stability.

## Methods

### Generation of yeast strains

Introduction of plasmids, gene deletions, and C-terminal fluorescent tagging was done by using lithium acetate-based yeast transformation^6^. All genomic manipulations were confirmed via plating on SC drop-out medium, PCR, and/or live-cell microscopy where applicable. Strains and plasmids used in this study are listed in Supplementary Table 1. Rad52-YFP and GFP-Tub1 under their endogenous promoters were expressed from a plasmid and secondary genomic locus, respectively.

### Microscopy

Live cell microscopy was done as described^16^. All experiments were done on logarithmic phase cells. For drug treatment, cells were treated with 0.03% MMS or 50 μg/mL zeocin for 1 hr. For combined MMS and 1,6-Hexanediol treatment, cells were treated with MMS for 1 hr, pelleted and washed with ddH_2_O, then re-suspended in SC drop-out medium with 5% 1,6-Hexanediol and digitonin [2 μg/mL] for 1 hr before imaging. Controls were treated similarly with digitonin. Small-budded S-phase cells in asynchronous cultures were subjected to live-cell confocal microscopy. Images were captured with a Leica DMI6000 SP8 LIGHTNING confocal microscope using a HC PL APO CS2 93x/1.30 glycerol objective. Pinhole was 155.3 μm and numerical aperture was 1.3. Images were deconvolved using Leica LIGHTNING deconvolution software and processed with Leica LAS software. Cells were maintained at 30°C throughout imaging. GFP was excited at 458 nm with a laser intensity of 28% and detected with a HyD hybrid detector set to 490 nm-523 nm. YFP was excited at 520 nm with an intensity of 3% and detected with a HyD hybrid detector set to 525-661 nm. For observation of liquid droplet behaviours, z-stack time-lapses were taken for 5-10 min. For Fig. 1d, n = 6 independent experiments. Each one of these 6 replicates consisted of the analysis of 30 cells that together provide ≥ 168 droplets. Cells were counted once. Maximum intensity projections are shown but all findings were confirmed on a single-plane. Particle velocities were measured using Imaris image analysis software.

### FLIP

Yeast strain KMY3426 was treated with 0.03% MMS and subjected to live cell confocal microscopy as described above. Images were acquired with a Leica SP8 LIGHTNING confocal microscope using an excitation wavelength of 488 nm, laser power of 0.2% and an HC PL APO CS2 63x/1.40 oil objective. GFP and YFP signals were detected and separated using HyD hybrid detectors set to 493-511 nm and 541-638 nm respectively. Pinhole was 95.5 μm and numerical aperture was 1.4. FLIP setup and subsequent analysis was done using Leica LAS software. For FLIP, S phase cells with Rad52-YFP foci were imaged 25 times before bleaching. A nuclear point outside the focus was bleached at 15% laser power for 25 ms and the cell was subsequently imaged. This bleaching/imaging process was repeated 25 times. The intensity of the focus was monitored throughout and normalized to the peak focus intensity.

### Protein isolation and immunoblotting

Briefly, 2.0×10^7^ cells were pelleted and protein was isolated and immunoblotting was performed as described^6^. Rad53 and Rad52 were detected using anti-Rad53 (Abcam-ab104232) and anti-Rad52 (gift from B. Pfander) antibodies. Chemiluminescence was captured using autoradiography and VersaDoc imager to ensure ideal exposure in linear range.

### Expression and purification of Rad52, *Δ307*, and RPA

Rad52 harbouring a C-terminal hexahistidine tag was expressed from plasmid pET11d, a kind gift from Dr. Lumir Krejci^23^. The Δ307 mutant was cloned from this template using PCR and standard methods and verified by sequencing. Rad52-His6 was expressed in *E.coli* BL21 Star pRARE. Luria broth (LB) cultures supplemented with 100 μg/mL ampicillin and 25 μg/mL chloramphenicol were inoculated with a single colony from a freshly transformed plate, grown overnight at 37°C, and diluted 100-fold into fresh LB containing 100 μg/mL ampicillin, 25 μg/mL chloramphenicol, and incubated at 37°C until OD600 0.6. Cultures were subsequently incubated at 16°C to an OD_600_ ~ 0.8 - 1. Expression of Rad52-His6 was induced by the addition of 0.1 mM isopropyl-D-thiogalactopyranoside and cultures were incubated at 16°C for 21 - 24 hr. Purification of Rad52-His6 was performed as described^23^. The purified protein was dialyzed against storage buffer (50 mM Tris-Cl pH 7.5, 100 mM KCl, 10% glycerol, 1 mM EDTA). As an alternative strategy, gel filtration through a Superdex 200 Increase 10/300 GL (GE Healthcare) column was used instead of hydroxyapatite chromatography^23^. The Δ307-His6 protein^25^ was produced similarly, with the following exceptions: i) cultures were grown in Terrific broth (TB) and ii) the final step of the purification involved gel filtration through a Superdex 200 Increase 10/300 GL (GE Healthcare) column in storage buffer (50 mM Tris-Cl pH 7.5, 100 mM KCl, 10% glycerol, 1 mM EDTA), instead of hydroxyapatite purification. Aliquots of all proteins were snap-frozen in liquid N_2_ and stored at −80°C. The Rpa heterotrimer was expressed from the pET11d plasmid^26^. Rpa was expressed and purified from *E. coli* BL21 Star pRARE cultures grown in TB, as described^30^.

### *In vitro* droplet assembly

Purified Rad52 protein was concentrated using an Amicon Ultra-0.5 mL centrifugal filter and suspended in glycerol-free storage buffer (50 mM Tris-Cl pH 7.5, 100 mM KCl, 1 mM EDTA). RPA and ssDNA complexes were generated *in vitro* by incubating 10 μM of purified RPA and 100 μM of PAGE-purified Oligo dT_30_ (IDT) in RPA storage buffer (60 mM HEPES pH 7.5, 0.5% Hyoinositol, 0.02% Tween 20, 0.5 mM EDTA) for 30 min at room temperature as described^31^. 20 μL reactions were performed in an uncoated 384-well coverslip plate (MatTek). Protein and glycerol-free dialysis buffer (50 mM Tris-Cl pH 7.5, 100 mM KCl, 1 mM EDTA) and/or salt-free glycerol-free dialysis buffer (50 mM Tris-Cl pH 7.5, 1 mM EDTA) were added to indicated concentrations. Reactions were imaged immediately following assembly. Images were captured with a Nikon C2+ Confocal Microscope using a Plan-Apochromat TIRF 60x oil objective, numerical aperture of 1.4 and pinhole of 30.0 μm. Phase separation *in vitro* was confirmed under indicated conditions using three independent Rad52 protein preparations.

### CFD

In order to simulate the dynamics of liquid droplets, the following governing equations of mass and momentum are solved using ANSYS-Fluent:

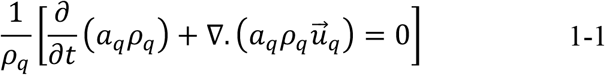

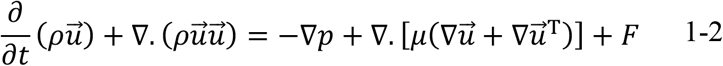

where, *a* represents the volume fraction of a phase, with 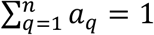, where *q* is a phase indicator (1 for the continuous phase, and 2 for the droplet), *ρ* is the volume average density and it is calculated by *ρ* = *a*_1_*ρ*_1_ + (1 – *a*_1_)*ρ*_2_, *u* is velocity, *t* is time, *p* is pressure, *μ* is viscosity and it is calculated by *μ* = *a*_1_*μ*_1_ + (1 – *a*_1_)*μ*_2_, and *F* is an external force. Continuum surface tension force (CSF) model is applied to model the surface tension force on the droplet interface. In this model, the pressure at the interphase is defined by *P*_2_ – *P*_1_ = 2*σ*/*R*, where *R* is the radius of the curvature (*κ*). The curvature is defined by the unit surface normal vector, where the normal vector is defined as the gradient of the volume fraction of the secondary phase, *n* = ∇*a_q_*: 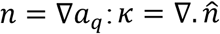, where 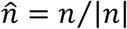. The surface tension force is *F* = 2*σρκ*∇*a*/(*ρ*_1_ + *ρ*_2_). This force is then inserted in equations above, which are solved numerically. Pressure and velocity are coupled with SIMPLE method. Pressure and momentum discretization methods are PRESTO and Second Order Upwind, respectively. Time is discretized by Second Order Implicit method.

Geo-Reconstruction method is applied for the volume fraction discretization. Inlet velocity is calculated as U = A cos ωt, where ω = 2 π f, A is average velocity, ω is angular frequency and f is frequency.

### Statistical analysis

Mann-Whitney U and *χ*^2^ square tests were used to compare non-normally distributed and categorical datasets respectively using GraphPad Prism 7.

### Data availability

All data are in the paper and supplementary information.

### Code availability

Codes are available on reasonable request.

## Acknowledgments

We thank the D. Durocher, G. Brown, and R. Hakem labs for reagents and discussions. We thank B. Pfander and U. Mortensen for reagents and strains. R.O. is supported by a Doctoral Scholarship from the Natural Sciences and Engineering Research Council of Canada (NSERC). The work was supported by a Canadian Institutes of Health Research (CIHR) grant (156297) to H.W., who holds the Canada Research Chair in Mechanisms of Genome Instability (950-231487). The work was mainly supported by funding from the CIHR (388041, 399687) and the Ontario Ministry of Research and Innovation (MRI-ERA; ER13-09-111) to K.M., who holds the Canada Research Chair in Spatial Genome Organization (CRC; 950-230661).

## Author contributions

R.O. and K.M. conceived the study and wrote the paper. Text editing by all co-authors. R.O. was the lead on all experiments except for the bacterial expression system (H.W. with E.Y.W.T) and *in vitro* droplet assays (R.O. and H.O.L.). M.M. executed CFD analyses under the guidance of N.A. and K.M. R.H. contributed to immunoblotting, strain making, and microscopy.

## Competing interests

Authors declare no competing interests.

## Additional Information

**Supplementary information** is available for this paper.

**Correspondence and requests for materials** should be addressed to K.M.

