## Supplementary Figures for "DNA repair by Rad52 liquid droplets"

### Supplementary Information

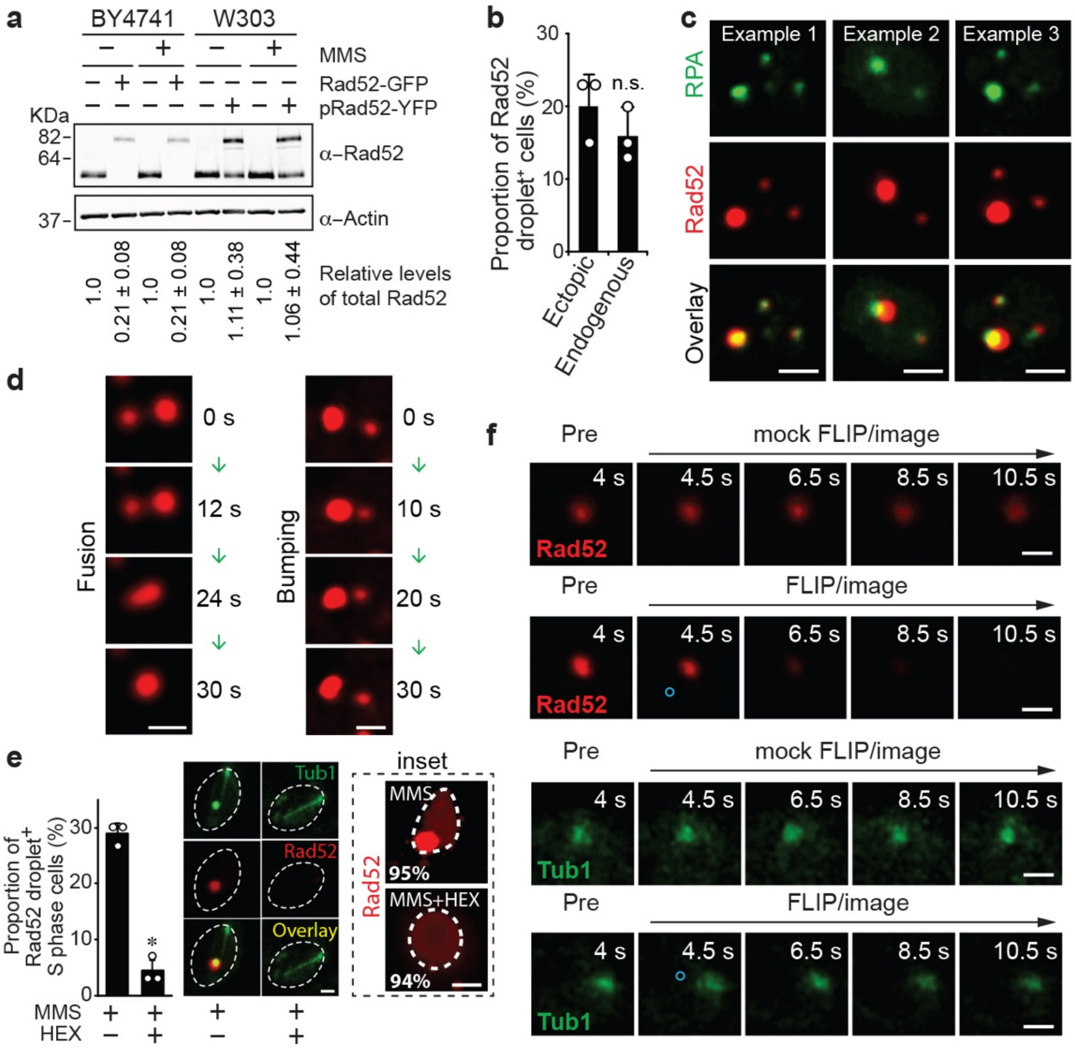


**Supplementary Figure 1 | DSB-indicating Rad52 foci behave like liquid droplets *in vivo.* a-b**, Plasmid-borne expression of fluorescently labelled Rad52, but not endogenously tagged Rad52, preserves total Rad52 protein levels (**a**) and MMS-induced Rad52 focus formation is similar when Rad52 is endogenously or exogenously tagged (**b**). Quantifications and standard deviations of total Rad52 protein levels in each lane normalized to its respective untagged control from three independent experiments are presented below the blots. **c**, Live cell microscopy shows that all Rad52 foci observed associate with an RPA focus (n=3, 251 foci). **d**, Zeocin-induced Rad52 droplets exhibit liquid droplet behaviours (scale bar, 0.5 μM). **e**, 1,6-Hexanediol disrupts Rad52 liquid droplets (n=3, 120 cells). Inset: 1,6-hexanediol disrupts MMS-induced Rad52 foci without altering the percentage of Rad52 fluorescence that is associated with the nucleus (dashed outline). (scale bar, 0.5 μm). **f**, Representative images of Rad52-YFP and GFP-Tub1 signal loss during FLIP experiments. Scale bars, 1 μM. Quantifications represent the mean ± s.d.; *$P < 0.0001$ in $\chi^2$ test.

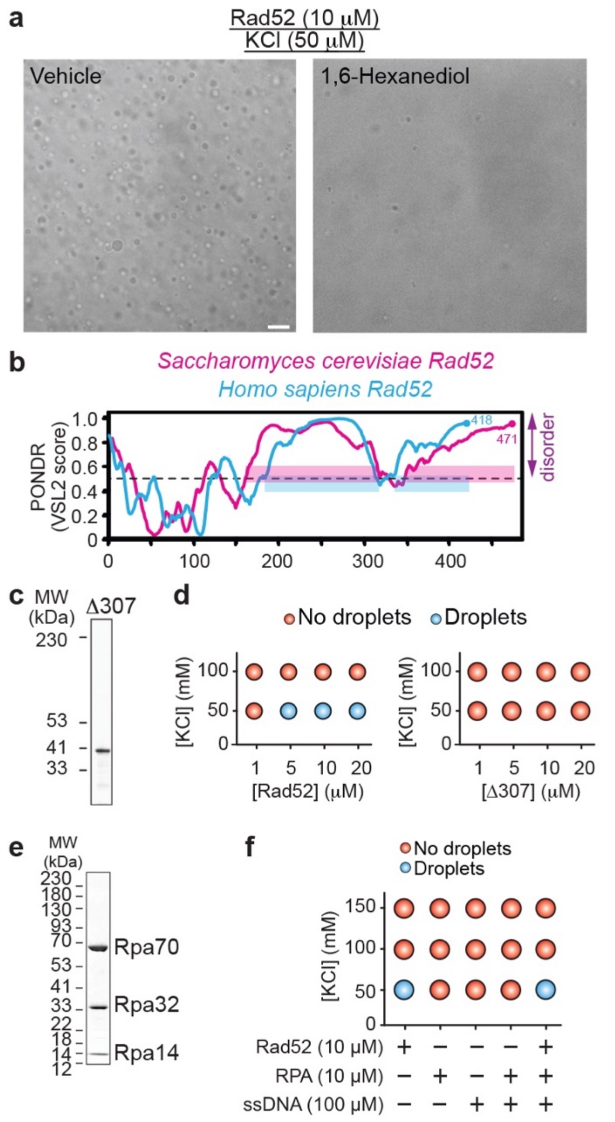


**Supplementary Figure 2 | Factors that influence Rad52 phase-separation in *vitro*. a**, Addition of 1,6-Hexanediol disrupts pre-formed Rad52 liquid droplets *in vitro*. Scale bar, 10 μm. **b**, Predictor of Natural Disordered Regions (PONDR) reveals that yeast and human Rad52 display similar disorder profiles across their amino acid sequences. **c-d**, Purified Rad52 C-terminal truncation Δ*307* fails to form droplets *in vitro*. **e-f**, Addition of RPA in complex with ssDNA does not influence Rad52 phase separation at varying salt concentrations.

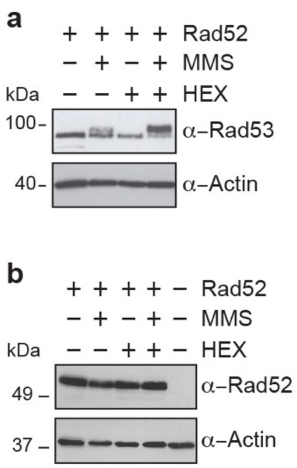


**Supplementary Figure 3| Phase-separation disruptor 1,6-Hexanediol hyper-activates the DNA damage checkpoint only in the presence of DNA damage induction. a-b**, Treatment with 1,6-Hexanediol (HEX) leads to hyperactivation of the Rad53 DNA damage checkpoint protein only in the prescence of MMS (**a**) without altering cellular Rad52 protein levels (**b**).

a

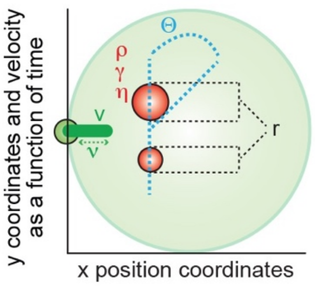


b

| Physical Properties | Calculated (C) or Tested (T) | Parameter values |
|---|---|---|
| Droplet size ratio (r) | C/T | **0.46** |
| Droplet density (ρ) | C/T | **1020** µg/µL |
| Droplet Surface tension (γ) | T | 1, 10, 100 µN/m |
| Droplet viscosity (η) | T | **0.005**, 0.5, 10 Pa s |
| Angle range to y-axis (Θ) | C/T | 0, 90 degrees |
| Intermitant MT velocity (v) | C/T | 20, 40 nm/s |
| Frequency of velocity (ν) | T | 10, 100, 1000 Hz |

**Supplementary Figure 4│ Parameters of CFD simulations of Rad52 liquid droplet fusions. a-b**, Schematic (**a**) and list (**b**) illustrating key parameters incorporated in the CFD simulations. Parameter values underlined are compatible with fusion while those bolded result in the most efficient fusion kinetics.

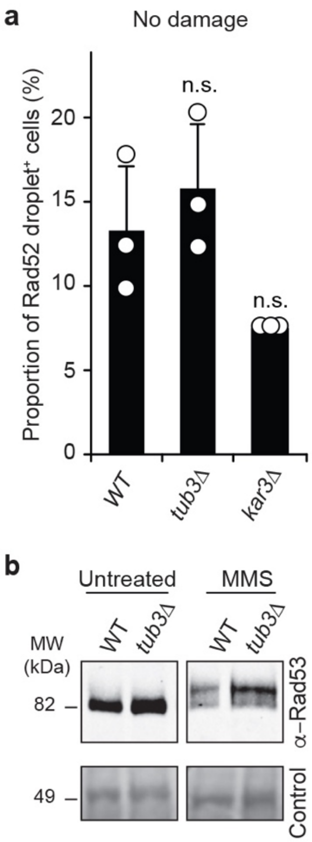


**Supplementary Figure 5│ Endogenous levels of Rad52 droplets. a**, Quantification of Rad52-YFP droplets via live-cell microscopy. Quantifications represent the mean ± s.d.; $\chi^2$ test was used. **b**, Longer running gel electrophoresis followed by western blotting shows that Tub3 deletion induces a higher migrating band indicating Rad53 hyper-phosphorylation. Ponceau serves as loading control.

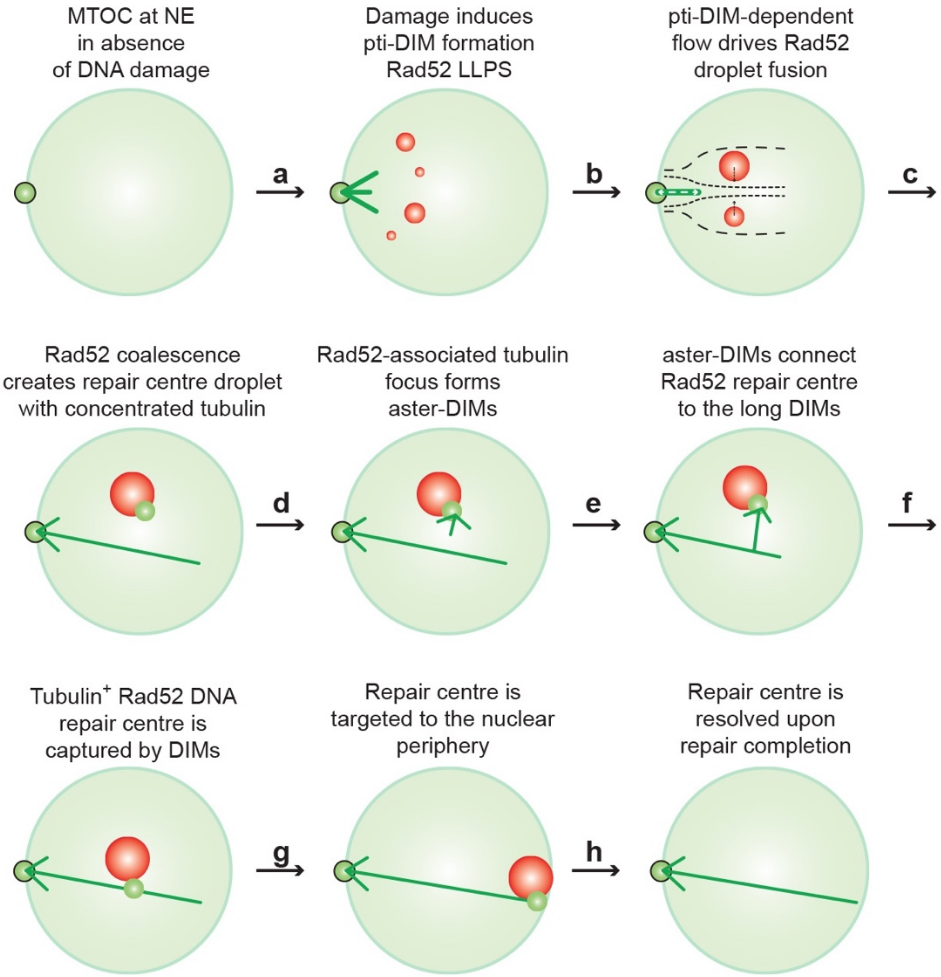


**Supplementary Figure 6 | Model**. Upon the induction of multiple DNA breaks, the evolutionarily conserved Rad52 protein assembles liquid droplets (red spheres) at the sites of DNA damage across the nucleus (large faint green sphere) (**a**). Novel intranuclear filaments termed pti-DIMs (short green sticks) also emerge from the nuclear envelope-embedded MTOC (**a**). Extension-shortening motions of pti-DIMs can then create a nucleoplasmic flow driving the fusion of the separate Rad52 droplets into a DNA repair centre droplet (**b**, **c**). The repair centre droplet internally concentrates tubulin and projects novel aster-like nuclear filaments termed aster-DIMs (**c**, **d**). One of the latter filaments matures and allows the Rad52 repair centre droplet to be captured by a long DIM (**e**, **f**) for mobilization to the nuclear periphery for repair (**g**, **h**). MTOC, microtubule organizing centre; NE, nuclear envelope; pti-DIMs, petite DNA damage-inducible intranuclear microtubule filaments; LLPS, liquid-liquid phase separation; aster-DIMs, aster-like DNA damage-inducible intranuclear microtubule filaments.

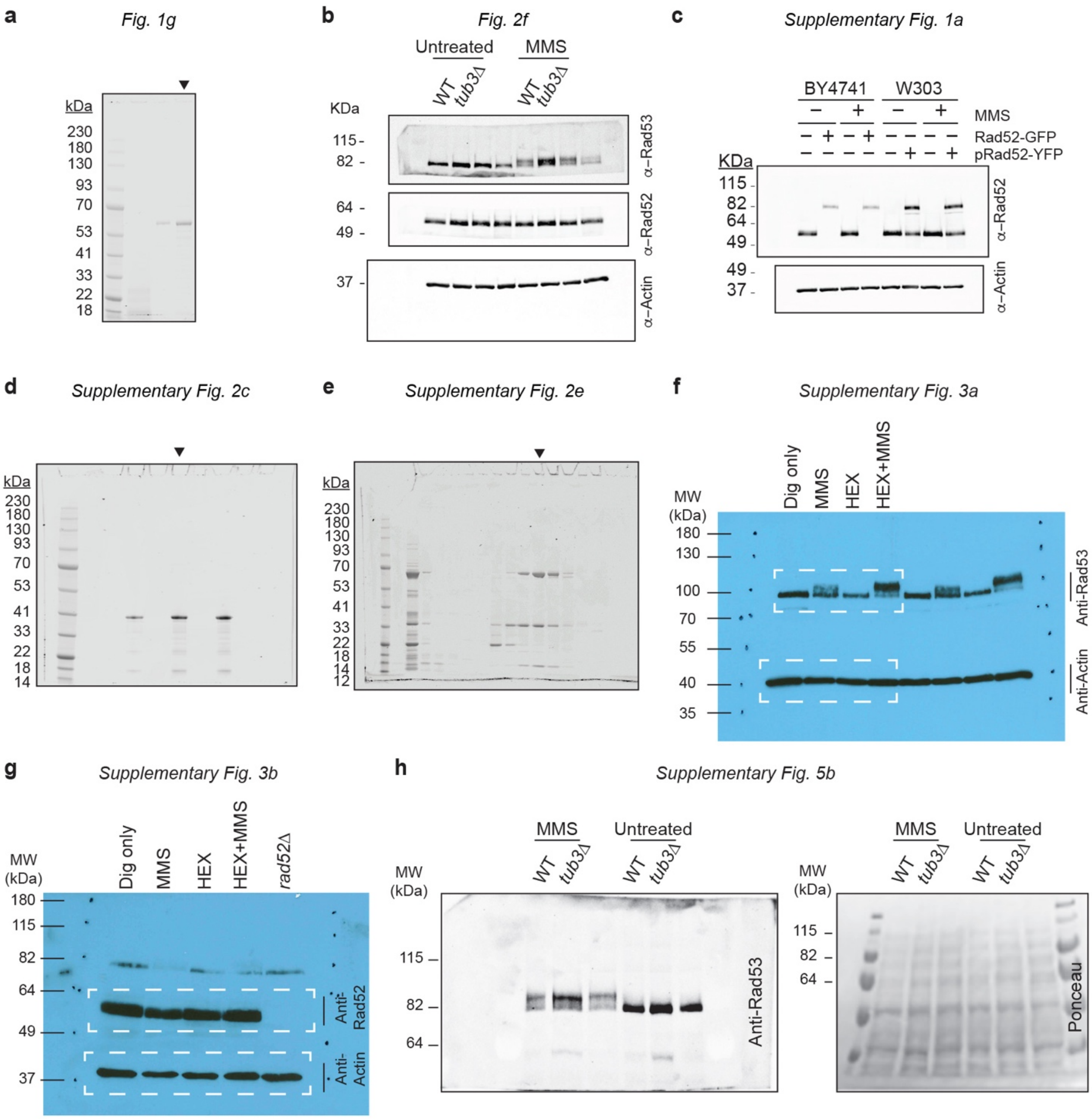


**Supplementary Figure 7│Uncropped SDS-PAGE gels and immunoblots**. Uncropped gels and immunoblots related to indicated main and supplementary figure panels.

**Supplementary Table 1 | List of S. cerevisiae strains and plasmids used in this study.**

| Reagent | Genotype | Source |
|---|---|---|
| **KMY 2559** | W303 MATa ade2-1 trp1-1 can1-100 leu2-3,112 his3-11,15 ura3-1 GAL+ psi+ ssd1-d2 RAD5+ | This study |
| **KMY 372** | BY4741 MATa his3Δ1 leu2Δ0 met15Δ0 ura3Δ0 | This study |
| **KMY 3426** | W303 MATa ade2-1 trp1-1 can1-100 leu2-3 URA3 GFP-Tub1-HIS3 nup49-GFP-KanMX pKM199-(Rad52-YFP-TRP1) | This study |
| **KMY 3557** | BY4741 Rfa1-GFP-HIS3+ pKM201 (Rad52-YFP-LEU2) | This study |
| **KMY 3542** | BY4741 Rad52-GFP-HIS3 | Kind gift from Brown lab |
| **KMY 3581** | KMY 3426 tub3Δ::HphR | This study |
| **KMY 3459** | KMY 3426 kar3Δ::HphR | This study |
| **KMY 3587** | KMY 3426 htz1Δ::HphR | This study |
| **pKM199** | Rad52-YFP-TRP1 | Kind gift from Durocher lab |
| **pKM 201** | Rad52-YFP-LEU2 | This study |
| **pKM397** | (His)6-RAD52(N+M)::pET11d | Kind gift from Krejci lab |
| **pKM 412** | (His)6-RAD52Δ307 (N+M)::pET11d | This study |
| **pKM 403** | p11d-sctRPA | Addgene |
