## Supplementary material for "DNA repair by Rad52 liquid droplets": Movie Legends

### DESCRIPTION OF ADDITIONAL SUPPLEMENTARY MATERIALS

**Supplementary Movie 1:** Live-cell time-lapse confocal microscopy showing multiple Rad52 foci (Rad52-YFP) engaging in dripping behaviour in a representative cell that was treated with 0.03% MMS to induce DNA damage. Scale bar, 1 µm.

**Supplementary Movie 2:** Live-cell time-lapse confocal microscopy reveals that Rad52 foci (Rad52-YFP) can fuse with each other in a representative cell that was treated with 0.03% MMS to induce DNA damage. Two fusion events are seen on the right. Scale bar, 1 µm.

**Supplementary Movie 3:** Live-cell time-lapse confocal microscopy showing multiple smaller Rad52 foci (Rad52-YFP) bumping into one larger central droplet in a yeast cell that was treated with 0.03% MMS to induce DNA damage. Scale bar, 1 µm.

**Supplementary Movie 4:** Live-cell time-lapse confocal microscopy from a FLIP experiment. Rad52 droplet (Rad52-YFP) intensity rapidly decreases upon photobleaching of a nucleoplasmic point outside the Rad52 focus. Yeast cells were treated with 0.03% MMS to induce DNA damage.

**Supplementary Movie 5:** Light microscopy showing the fusion of Rad52 droplets assembled *in vitro*.

**Supplementary Movie 6:** Live-cell time-lapse confocal microscopy revealing dynamic pti-DIMs inside the nucleus. Cells were treated with 0.03% MMS to induce DNA damage. GFP-Tub1 (α-Tubulin) and Nup49-GFP (nuclear pore complexes) are shown in cyan. Scale bar, 1 µm.

**Supplementary Movie 7:** Time-lapse from Supplementary Movie 6 overlaid with the corresponding Rad52 droplet signal (Rad52-YFP shown in magenta). Scale bar, 1 µm.

**Supplementary Movie 8:** Representative computational flow dynamics simulation in which pti-DIM extension-shortening cycles resulted in the fusion of two Rad52 droplets. The line connecting the droplets in their initial position was perpendicular to the velocity exerted by the pti-DIMs at the centre of the Y-axis. The parameters used in this simulation were $r = 046$, $\rho = 1014$ $Kg/m^3$, $\sigma = 1.4$ uN/m, $\mu = 0.005$, $\Theta = 90°$, $u = 41$ nm/s, and $f = 1000$ Hz as shown in Supplementary Figure 2.

**Supplementary Movie 9:** Representative computational flow dynamics simulation in which pti-DIM extension-shortening cycles failed to result in the fusion of two Rad52 droplets. The line connecting the droplets in their initial position was rotated clockwise by 20° compared to the position in Supplementary Movie 8, thereby changing Θ to 110°. Other simulation parameters were identical to those used in that movie.

**Supplementary Movie 10:** Live-cell time-lapse confocal microscopy showing a large Rad52 droplet with an internally concentrated tubulin focus that is captured by and travels along a stable DIM. Cells were treated with 0.03% MMS to induce DNA damage. GFP-Tub1 (tubulin) and Nup49-GFP (nuclear envelope) are shown in cyan and Rad52-YFP in magenta. Scale bar, 1 µm.
